# Retrospective serological analysis reveals presence of the emerging lagovirus RHDV2 in Australian wild rabbits at least six months prior to its first detection

**DOI:** 10.1101/613158

**Authors:** Tanja Strive, Melissa Piper, Nina Huang, Roslyn Mourant, John Kovaliski, Lorenzo Capucci, Tarnya E Cox, Ina Smith

## Abstract

The lagovirus Rabbit Haemorrhagic Disease Virus (RHDV) has been circulating in Australia since the mid-1990s when it was deliberately released to control overabundant rabbit populations. In recent years, the viral diversity of different RHDVs in Australia has increased, and currently four different types of RHDV are known to be circulating. To allow for ongoing epidemiological studies and impact assessments of these viruses on Australian wild rabbit populations, it is essential that serological tools are updated. To this end, Reference sera were produced against all four virulent RHDVs (including RHDV2) known to be present in Australia and tested in a series of available immunological assays originally developed for the prototype RHDV, to assess patterns of cross reactivity and the usefulness of these assays to detect lagovirus antibodies, either in a generic or specific manner. Enzyme Linked Immuno Sorbent Assays (ELISAs) developed to detect antibody isotypes IgM, IgA and IgG were sufficiently cross reactive to detect antibodies raised against all four virulent lagoviruses. For the more specific detection of antibodies to the antigenically more different RHDV2, a competition ELISA was adapted using RHDV2 specific monoclonal antibodies in combination with Australian viral antigen. Archival serum banks from a long term rabbit monitoring site where rabbits were sampled quarterly over a period of six years were re-screened using this assay, and revealed serological evidence for the arrival of RHDV2 in this population at least six months prior to its initial detection in Australia in a deceased rabbit in May 2015. The serological methods and reference reagents described here will provide valuable tools to study presence, prevalence and impact of RHDV2 on Australian rabbit populations; however the discrimination of different antigenic variants of RHDVs as well as mixed infections at the serological level remains challenging.

## Introduction

Rabbit Haemorrhagic Disease Virus (RHDV, sometimes also referred to as RHDV1 or GI.1c, according to a new proposed nomenclature (Le Pendu et al., 2017)), belongs to the genus lagovirus within the family *caliciviridae*. RHDV was released in Australia as a biological control agent for introduced wild rabbits, a devastating agricultural and environmental pest species in this country (Cooke and Fenner, 2002). While initially very effective in reducing rabbit populations across large parts of the continent (Saunders et al., 2002, Mutze et al., 1998), RHDV was not effective in the more temperate areas of South Eastern Australia. This lack of effectiveness was likely due to the presence of endemic, non-pathogenic caliciviruses (Rabbit calicivirus Australia 1, also termed RCV-A1, or GI.4) that can provide transient and partial immunological cross protection to RHDV (Strive et al., 2013, Liu et al., 2014), thereby reducing both case fatality and infection rates (Cooke et al., 2018). In addition to the impeding effects of RCV-A1, rabbit populations have been recovering in recent years (Mutze et al., 2014), and developing genetic resistance to RHDV has been reported in some Australian rabbit populations (Elsworth et al., 2012, Nystroem et al., 2011). In an attempt to ‘boost’ rabbit biocontrol in Australia and to maintain the substantial economic and environmental gains made by the long term suppression of rabbit populations by RHDV (Pedler et al., 2016, Cooke, 2013), an additional strain of RHDV was released nationwide in Australia in March 2017 (Hall et al., 2018, Strive and Cox, 2019). This strain termed RHDVa-K5 is a naturally occurring antigenic variant of RHDV from Korea (Oem et al., 2009) which was shown experimentally to be more effective in infecting rabbits from a genetically resistant rabbit population (Elsworth et al., 2012) and in overcoming partial protection conveyed by the benign RCV-A1 (Cox et al., 2013). These antigenic variants of RHDV, referred to as RHDVa (GI.1a, or RHDV1a), were first reported in the late 1990s (Capucci et al., 1998), and although they exhibit antigenic differences they are considered to be of the same serotype (Lavazza and Capucci, 2016).

Prior to the release of RHDVa-K5, the incursions of two additional RHDV variants were reported in Australia. The first incursion was another variant RHDVa (GI.1a) strain in the greater Sydney area that most closely resembled a Chinese isolate (Wang et al., 2012) that appeared in early 2014. This virus, termed RHDVa-Aus, caused a number of recorded outbreaks mostly in domestic rabbit farms, however, its distribution appeared geographically limited to the east and north east of New South Wales (Mahar et al., 2018b). In May 2015, the incursion of a second exotic virus, the recently emerged RHDV2 (GI.2) was reported in Australia (Hall et al., 2015). RHDV2 is a new lagovirus that was first reported in Europe in 2010 (Le Gall-Recule et al., 2011, Dalton et al., 2012). It is not only genetically distinct from RHDV and RHDVa but, unlike RHDV and RHDVa, is also able to cause highly fatal disease in very young rabbits (Neimanis et al., 2018a, Dalton et al., 2014) and capable of fatally infecting several species of hares, making it the only known lagovirus that does not exhibit strict species specificity (Camarda et al., 2014, Hall et al., 2017, Le Gall-Recule et al., 2017, Neimanis et al., 2018b, Puggioni et al., 2013, Velarde et al., 2017). Since its arrival in Australia, RHDV2 has become the dominant strain circulating in the field, seemingly replacing older RHDV1 strains and accounting for the majority of reported cases in wild and domestic rabbits (Mahar et al., 2018a).

With the increasing diversity of co-circulating RHDVs both in Australia and Europe it was essential to update the diagnostic tools available, to allow for specific identification of strains and to measure the impact they were having on wild rabbit (and hare) populations. Due to the genetic variability, the development of specific molecular diagnostics was comparably straightforward, and accordingly several tailored approaches have been described in both Europe and Australia (Carvalho et al., 2017, Duarte et al., 2015, Hall et al., 2018, Le Gall-Recule et al., 2017). In contrast, the development of specific serological tools is more challenging. Although strain specific epitopes have been described for lagoviruses and specific monoclonal antibodies have been developed (Capucci et al., 1996a, Capucci et al., 1998, Capucci et al., 1995, Dalton et al., 2018, Kong et al., 2016, Liu et al., 2012b), discriminatory serology remains difficult, in particular for closely related RHDVs, due to the large number of shared epitopes (Lavazza and Capucci, 2016). The infection history of populations can often only be inferred based on reactivity patterns (Barcena et al., 2015, Velarde et al., 2016).

In Australia, the development of specific serological tools to discriminate between different lagoviruses was of particular importance. Here, RHDV2 was actively circulating at the time the new strain RHDVa-K5 was released nationwide (Strive and Cox, 2019) and specific serological tools were needed to investigate the respective impacts and potential interactions the two strains had on wild Australian rabbits at a population level. Furthermore, in the years leading up to the national release of RHDVa-K5, extensive serological monitoring of several Australian rabbit populations was carried out. Screening these sample banks with a serological assay specific to RHDV2 might allow a more accurate determination of the exact time point this virus entered into Australia and started to circulate in wild rabbit populations before it was detected in May 2015 (Hall et al., 2015).

This study describes the production of reference sera raised against all virulent RHDV strains known to be circulating in Australia for the purpose of assessing the cross-reactivity patterns in a panel of existing serological assays used to infer disease dynamics of RHDV in rabbit populations. It further describes the adaptation of a European RHDV2 cELISA to the Australian strain of RHDV2, and the modification of IgM and IgA isotype ELISAs for the improved detection of RHDV2 antibodies. The RHDV2 cELISA was then applied to retrospectively analyse a long term field monitoring site to estimate the time of arrival on RHDV2 at this site.

## Material and methods

### Ethics approval

All work involving animals was approved by the CSIRO Ecosystem Sciences Animal Ethics Committee (ESAEC #10-13, #11-01, #13-01, #13-10, #DOMRAB) and the Orange Animal Ethics Committee (ORA11/14/001, ORA14/17/001) and carried out in accordance with the Australian code for the care and use of animals for scientific purposes.

### Production of antigen, virus inoculum and experimental vaccine

Two five week old New Zealand white rabbits were infected with 0.5 ml of a 2% clarified liver homogenate of the first RHDV2 field isolate found in Australia (BlMt-1, Gen Bank# KT280060) (Hall et al., 2015). Rabbits were monitored twice daily for signs of terminal rabbit haemorrhagic disease (RHD), which was defined as 10% weight loss within a 24 hour period, no resistance to handling or lateral recumbency, or hypothermia following a fever episode, often in combination with lethargy. Both rabbits experienced a peracute form of RHD, one was found displaying signs of terminal RHD at 66 hours post infection (h.p.i.) and was euthanized, the second was found dead 90 h.p.i. with no prior signs of terminal RHD detected.

The liver of one of the RHDV2 infected rabbits was used for the production of an experimental vaccine, according to methods published previously (Lavazza and Capucci, 2016). Briefly, a 20% w/v liver homogenate (containing approximately 3 × 10^8^ capsid gene copies/ml) was prepared in sterile PBS and clarified by centrifugation for 20 min at 2,000 g. Chloroform was added to the supernatant to a final concentration of 2% (v/v) and incubated at 4°C overnight, followed by a second clarification at 10,000g and 4°C. Part of the clarified supernatant was removed, mixed 1:1 with glycerol and stored at −80°C as RHDV2 inoculum for subsequent infections (0.5 ml/rabbit). Formalin was added to the remaining supernatant to a final concentration of 0.8% v/v and incubated at room temperature overnight. A second inactivation step was performed by adding formalin to a total concentration of 1% v/v with overnight incubation at 4°C. This preparation was stored at 4°C until used. Immediately prior to use, the vaccine was brought to room temperature, mixed 1:1 with Addavax (Invivogen, San Diego, USA), and between 0.6 and 1.2 ml were injected subcutaneously in the scruff of the neck.

For the production of RHDV2 ELISA antigen, a 10% w/v homogenate of the liver of the second rabbit was clarified using two centrifugation steps (20 min at 3000g, followed by 30 min at 6000g, at 4°C), and then passed through a 0.8 µm filter. The resulting homogenate was mixed 1:1 with glycerol and stored at −80°C.

A commercially produced preparation of RHDVa-K5 was used as inoculum to produce hyperimmune sera, diluted to 10,000 RID_50_/ml (Elizabeth Macarthur Agricultural Institute, Menangle, Australia). RHDVa-Aus inoculum was prepared from a 2% of clarified liver homogenate of RHDVa-Aus (Ber-3, GenBank # KY628310) (Mahar et al., 2018b). The infectious dose of the RHDVa-Aus and the RHDV2 inoculum was not titrated.

### Production of reference sera

For the production of RHDVa hyperimmune sera, five-week old rabbits were infected with RHDVa-Aus (n=4) and RHDVa-K5 (n=5). Due to the age related innate resistance to lethal RHDV infection (Matthaei et al., 2014, Neave et al., 2018) it was expected that rabbits of this age group would not succumb to fatal RHDV infection, but survive and mount a strong antibody response. Rabbits were orally infected with 0.5 ml of virus inoculum using a 1 ml syringe. Rabbits were monitored twice daily for the first four days and then daily afterwards. A small (0.1-0.2 ml) blood sample was collected from the marginal ear vein at day 0, 7 and 14. Two rabbits in each group were sacrificed and bled at 14 dpi the remaining rabbits at 20 dpi (RHDVa-K5) and 22 dpi (RHDVa-Aus) (Table 1).

**Table 1:**
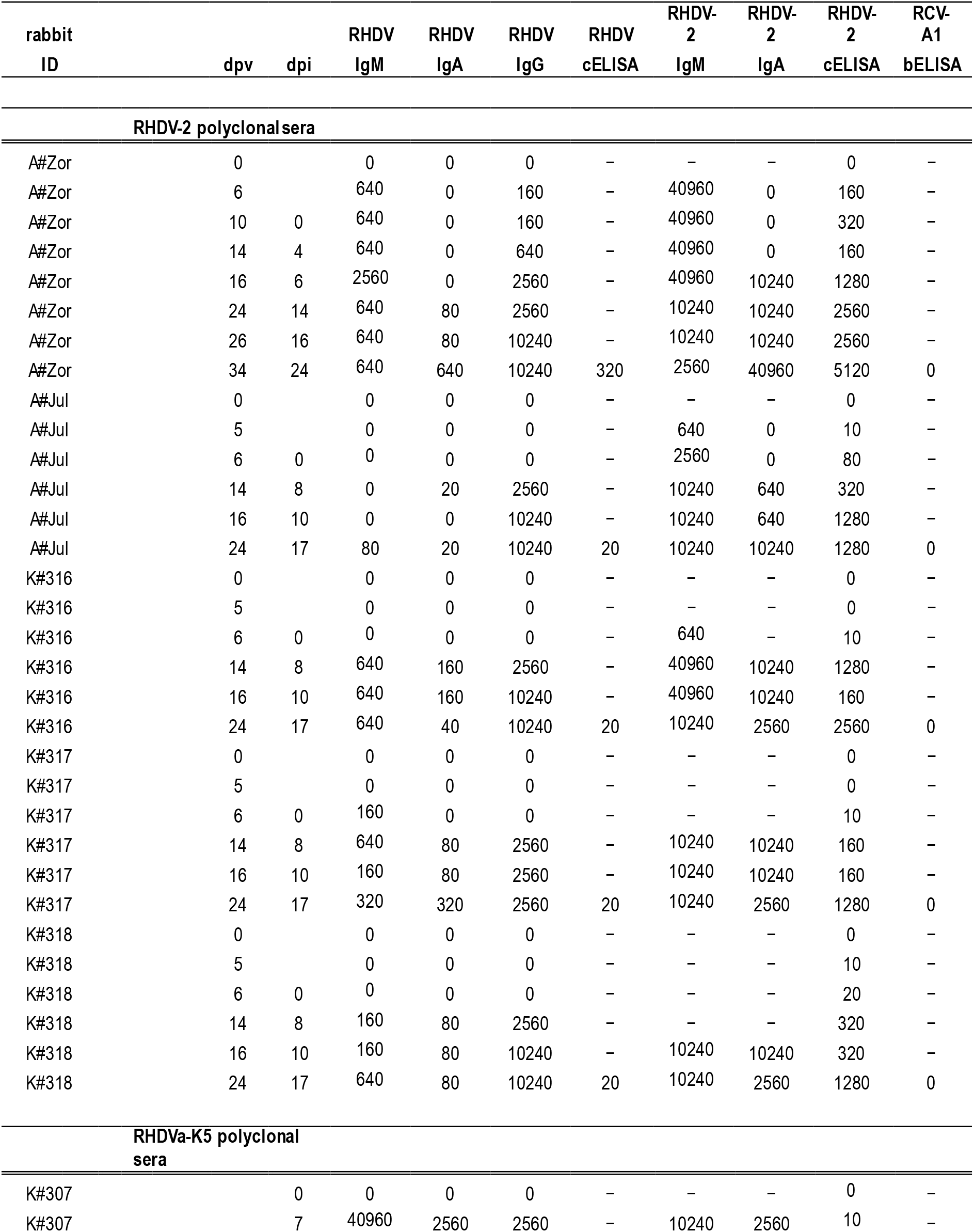
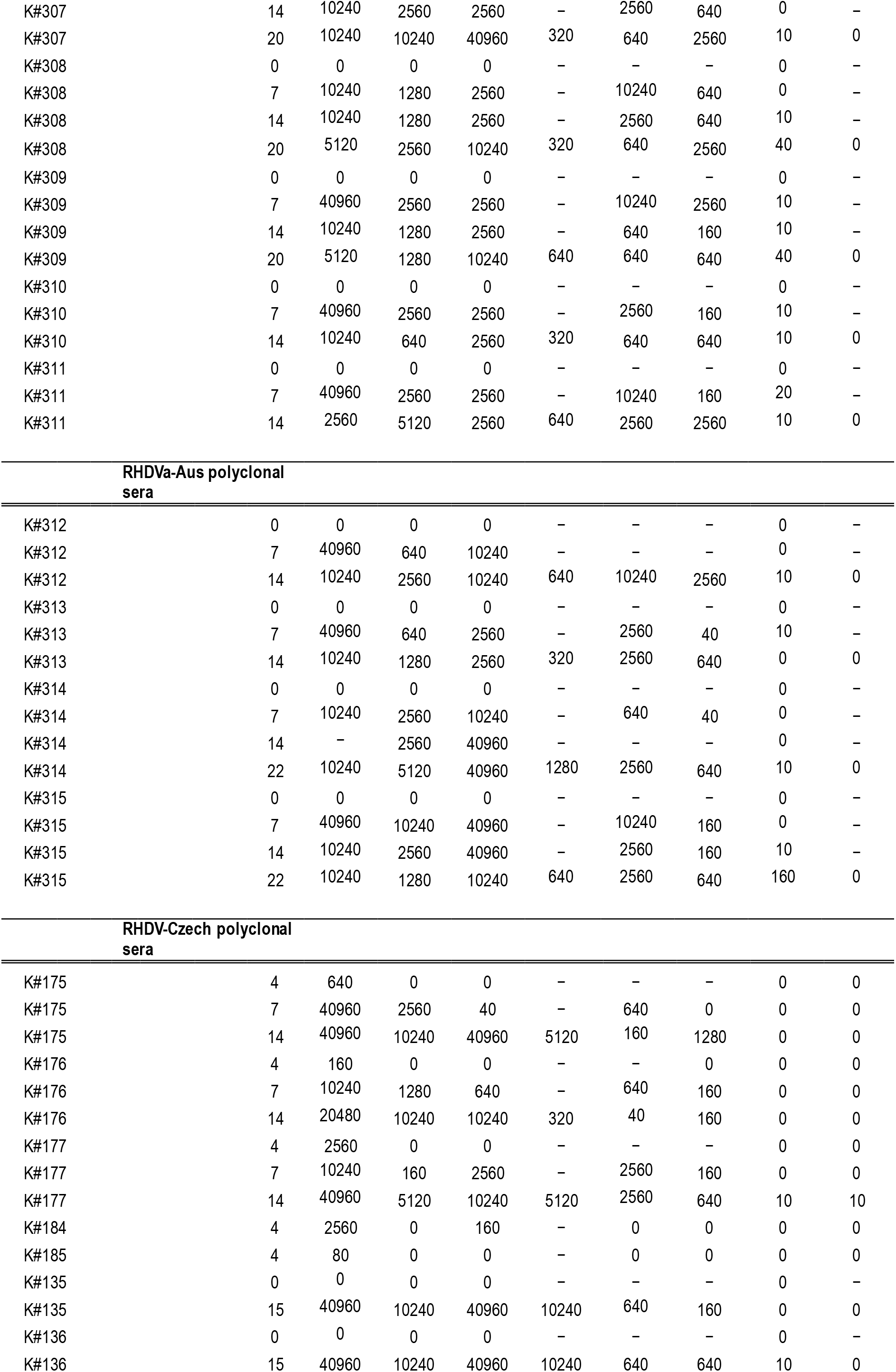
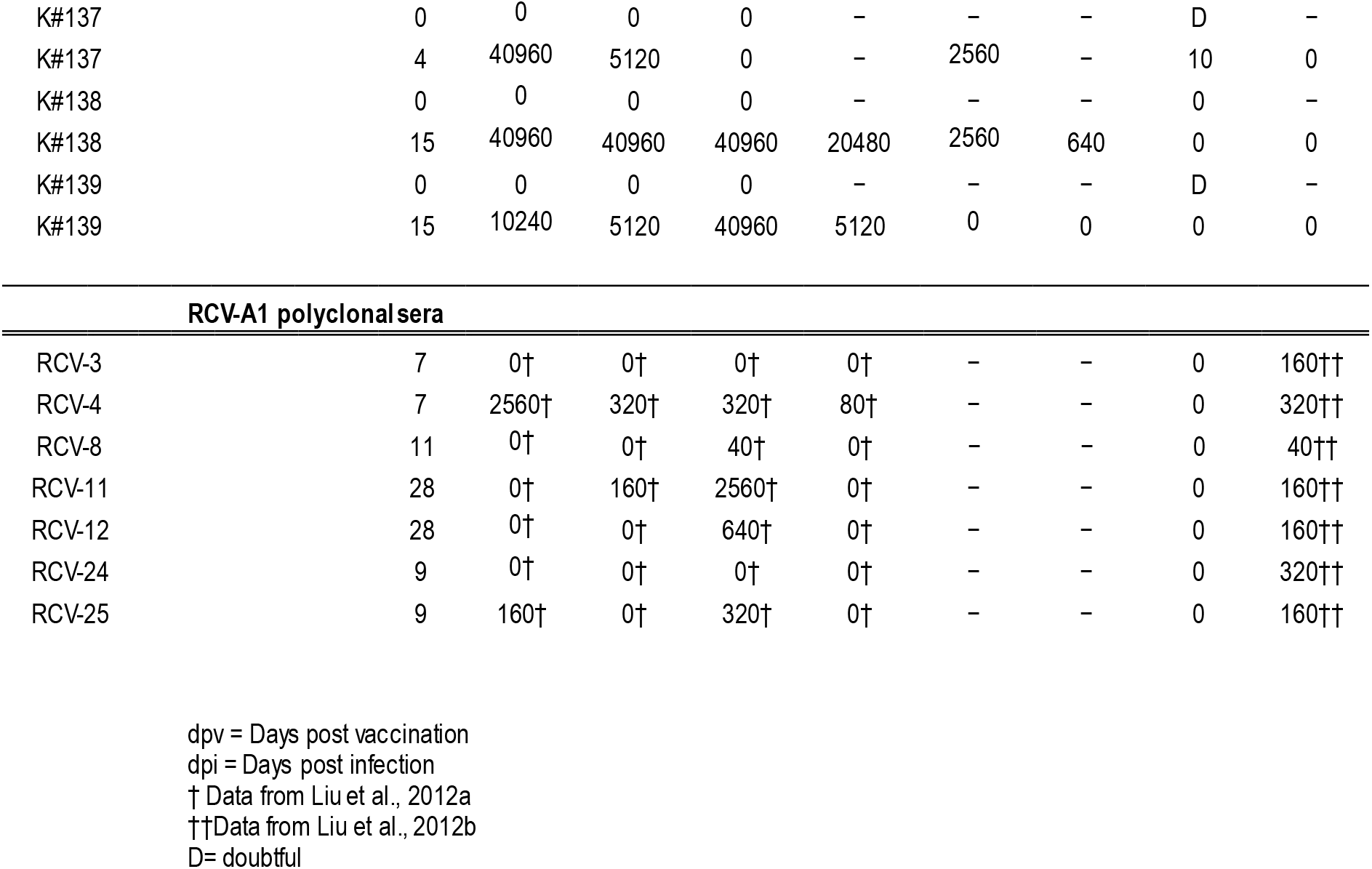
Serological reaction profiles in experimentally generated polyclonal sera against four types of RHDV and RCV-A1.

| rabbit ID | dpv | dpi | RHDV IgM | RHDV IgA | RHDV IgG | RHDV cELISA | RHDV-2 IgM | RHDV-2 IgA | RHDV-2 cELISA | RCV-A1 bELISA |
| --- | --- | --- | --- | --- | --- | --- | --- | --- | --- | --- |
| **RHDV-2 polyclonal sera** |  |  |  |  |  |  |  |  |  |  |
| A#Zor | 0 |  | 0 | 0 | 0 | - | - | - | 0 | - |
| A#Zor | 6 |  | 640 | 0 | 160 | - | 40960 | 0 | 160 | - |
| A#Zor | 10 | 0 | 640 | 0 | 160 | - | 40960 | 0 | 320 | - |
| A#Zor | 14 | 4 | 640 | 0 | 640 | - | 40960 | 0 | 160 | - |
| A#Zor | 16 | 6 | 2560 | 0 | 2560 | - | 40960 | 10240 | 1280 | - |
| A#Zor | 24 | 14 | 640 | 80 | 2560 | - | 10240 | 10240 | 2560 | - |
| A#Zor | 26 | 16 | 640 | 80 | 10240 | - | 10240 | 10240 | 2560 | - |
| A#Zor | 34 | 24 | 640 | 640 | 10240 | 320 | 2560 | 40960 | 5120 | 0 |
| A#Jul | 0 |  | 0 | 0 | 0 | - | - | - | 0 | - |
| A#Jul | 5 |  | 0 | 0 | 0 | - | 640 | 0 | 10 | - |
| A#Jul | 6 | 0 | 0 | 0 | 0 | - | 2560 | 0 | 80 | - |
| A#Jul | 14 | 8 | 0 | 20 | 2560 | - | 10240 | 640 | 320 | - |
| A#Jul | 16 | 10 | 0 | 0 | 10240 | - | 10240 | 640 | 1280 | - |
| A#Jul | 24 | 17 | 80 | 20 | 10240 | 20 | 10240 | 10240 | 1280 | 0 |
| K#316 | 0 |  | 0 | 0 | 0 | - | - | - | 0 | - |
| K#316 | 5 |  | 0 | 0 | 0 | - | - | - | 0 | - |
| K#316 | 6 | 0 | 0 | 0 | 0 | - | 640 | - | 10 | - |
| K#316 | 14 | 8 | 640 | 160 | 2560 | - | 40960 | 10240 | 1280 | - |
| K#316 | 16 | 10 | 640 | 160 | 10240 | - | 40960 | 10240 | 160 | - |
| K#316 | 24 | 17 | 640 | 40 | 10240 | 20 | 10240 | 2560 | 2560 | 0 |
| K#317 | 0 |  | 0 | 0 | 0 | - | - | - | 0 | - |
| K#317 | 5 |  | 0 | 0 | 0 | - | - | - | 0 | - |
| K#317 | 6 | 0 | 160 | 0 | 0 | - | - | - | 10 | - |
| K#317 | 14 | 8 | 640 | 80 | 2560 | - | 10240 | 10240 | 160 | - |
| K#317 | 16 | 10 | 160 | 80 | 2560 | - | 10240 | 10240 | 160 | - |
| K#317 | 24 | 17 | 320 | 320 | 2560 | 20 | 10240 | 2560 | 1280 | 0 |
| K#318 | 0 |  | 0 | 0 | 0 | - | - | - | 0 | - |
| K#318 | 5 |  | 0 | 0 | 0 | - | - | - | 10 | - |
| K#318 | 6 | 0 | 0 | 0 | 0 | - | - | - | 20 | - |
| K#318 | 14 | 8 | 160 | 80 | 2560 | - | - | - | 320 | - |
| K#318 | 16 | 10 | 160 | 80 | 10240 | - | 10240 | 10240 | 320 | - |
| K#318 | 24 | 17 | 640 | 80 | 10240 | 20 | 10240 | 2560 | 1280 | 0 |
| **RHDVa-K5 polyclonal sera** |  |  |  |  |  |  |  |  |  |  |
| K#307 |  | 0 | 0 | 0 | 0 | - | - | - | 0 | - |
| K#307 |  | 7 | 40960 | 2560 | 2560 | - | 10240 | 2560 | 10 | - |

|  |  |  |  |  |  |  |  |  |  |
| --- | --- | --- | --- | --- | --- | --- | --- | --- | --- |
| K#307 | 14 | 10240 | 2560 | 2560 | - | 2560 | 640 | 0 | - |
| K#307 | 20 | 10240 | 10240 | 40960 | 320 | 640 | 2560 | 10 | 0 |
| K#308 | 0 | 0 | 0 | 0 | - | - | - | 0 | - |
| K#308 | 7 | 10240 | 1280 | 2560 | - | 10240 | 640 | 0 | - |
| K#308 | 14 | 10240 | 1280 | 2560 | - | 2560 | 640 | 10 | - |
| K#308 | 20 | 5120 | 2560 | 10240 | 320 | 640 | 2560 | 40 | 0 |
| K#309 | 0 | 0 | 0 | 0 | - | - | - | 0 | - |
| K#309 | 7 | 40960 | 2560 | 2560 | - | 10240 | 2560 | 10 | - |
| K#309 | 14 | 10240 | 1280 | 2560 | - | 640 | 160 | 10 | - |
| K#309 | 20 | 5120 | 1280 | 10240 | 640 | 640 | 640 | 40 | 0 |
| K#310 | 0 | 0 | 0 | 0 | - | - | - | 0 | - |
| K#310 | 7 | 40960 | 2560 | 2560 | - | 2560 | 160 | 10 | - |
| K#310 | 14 | 10240 | 640 | 2560 | 320 | 640 | 640 | 10 | 0 |
| K#311 | 0 | 0 | 0 | 0 | - | - | - | 0 | - |
| K#311 | 7 | 40960 | 2560 | 2560 | - | 10240 | 160 | 20 | - |
| K#311 | 14 | 2560 | 5120 | 2560 | 640 | 2560 | 2560 | 10 | 0 |

**RHDVa-Aus polyclonal sera**
|  |  |  |  |  |  |  |  |  |  |
| --- | --- | --- | --- | --- | --- | --- | --- | --- | --- |
| K#312 | 0 | 0 | 0 | 0 | - | - | - | 0 | - |
| K#312 | 7 | 40960 | 640 | 10240 | - | - | - | 0 | - |
| K#312 | 14 | 10240 | 2560 | 10240 | 640 | 10240 | 2560 | 10 | 0 |
| K#313 | 0 | 0 | 0 | 0 | - | - | - | 0 | - |
| K#313 | 7 | 40960 | 640 | 2560 | - | 2560 | 40 | 10 | - |
| K#313 | 14 | 10240 | 1280 | 2560 | 320 | 2560 | 640 | 0 | 0 |
| K#314 | 0 | 0 | 0 | 0 | - | - | - | 0 | - |
| K#314 | 7 | 10240 | 2560 | 10240 | - | 640 | 40 | 0 | - |
| K#314 | 14 | - | 2560 | 40960 | - | - | - | 0 | - |
| K#314 | 22 | 10240 | 5120 | 40960 | 1280 | 2560 | 640 | 10 | 0 |
| K#315 | 0 | 0 | 0 | 0 | - | - | - | 0 | - |
| K#315 | 7 | 40960 | 10240 | 40960 | - | 10240 | 160 | 0 | - |
| K#315 | 14 | 10240 | 2560 | 40960 | - | 2560 | 160 | 10 | - |
| K#315 | 22 | 10240 | 1280 | 10240 | 640 | 2560 | 640 | 160 | 0 |

**RHDV-Czech polyclonal sera**
|  |  |  |  |  |  |  |  |  |  |
| --- | --- | --- | --- | --- | --- | --- | --- | --- | --- |
| K#175 | 4 | 640 | 0 | 0 | - | - | - | 0 | 0 |
| K#175 | 7 | 40960 | 2560 | 40 | - | 640 | 0 | 0 | 0 |
| K#175 | 14 | 40960 | 10240 | 40960 | 5120 | 160 | 1280 | 0 | 0 |
| K#176 | 4 | 160 | 0 | 0 | - | - | 0 | 0 | 0 |
| K#176 | 7 | 10240 | 1280 | 640 | - | 640 | 160 | 0 | 0 |
| K#176 | 14 | 20480 | 10240 | 10240 | 320 | 40 | 160 | 0 | 0 |
| K#177 | 4 | 2560 | 0 | 0 | - | - | - | 0 | 0 |
| K#177 | 7 | 10240 | 160 | 2560 | - | 2560 | 160 | 0 | 0 |
| K#177 | 14 | 40960 | 5120 | 10240 | 5120 | 2560 | 640 | 10 | 10 |
| K#184 | 4 | 2560 | 0 | 160 | - | 0 | 0 | 0 | 0 |
| K#185 | 4 | 80 | 0 | 0 | - | 0 | 0 | 0 | 0 |
| K#135 | 0 | 0 | 0 | 0 | - | - | - | 0 | - |
| K#135 | 15 | 40960 | 10240 | 40960 | 10240 | 640 | 160 | 0 | 0 |
| K#136 | 0 | 0 | 0 | 0 | - | - | - | 0 | - |
| K#136 | 15 | 40960 | 10240 | 40960 | 10240 | 640 | 640 | 10 | 0 |
| K#137 | 0 | 0 | 0 | 0 | - | - | - | D | - |
| K#137 | 4 | 40960 | 5120 | 0 | - | 2560 | - | 10 | 0 |
| K#138 | 0 | 0 | 0 | 0 | - | - | - | 0 | - |
| K#138 | 15 | 40960 | 40960 | 40960 | 20480 | 2560 | 640 | 0 | 0 |
| K#139 | 0 | 0 | 0 | 0 | - | - | - | D | - |
| K#139 | 15 | 10240 | 5120 | 40960 | 5120 | 0 | 0 | 0 | 0 |

| RCV-A1 polyclonal sera |  |  |  |  |  |  |  |  |  |
| --- | --- | --- | --- | --- | --- | --- | --- | --- | --- |
| RCV-3 | 7 | 0† | 0† | 0† | 0† | - | - | 0 | 160†† |
| RCV-4 | 7 | 2560† | 320† | 320† | 80† | - | - | 0 | 320†† |
| RCV-8 | 11 | 0† | 0† | 40† | 0† | - | - | 0 | 40†† |
| RCV-11 | 28 | 0† | 160† | 2560† | 0† | - | - | 0 | 160†† |
| RCV-12 | 28 | 0† | 0† | 640† | 0† | - | - | 0 | 160†† |
| RCV-24 | 9 | 0† | 0† | 0† | 0† | - | - | 0 | 320†† |
| RCV-25 | 9 | 160† | 0† | 320† | 0† | - | - | 0 | 160†† |

dpv = Days post vaccination dpi = Days post infection † Data from Liu et al., 2012a †† Data from Liu et al., 2012b D= doubtful

Archival serum samples from five rabbits collected at various time points were used as RHDV hyperimmune reference sera. In this previous study (T. Strive, unpublished), three 5 week old New Zealand white rabbits were infected orally with 500 ID_50_ of a commercial RHDV preparation (RHDV v351, Elizabeth Macarthur Agricultural Institute, Menangle, Australia), blood samples were collected at 7 dpi and a final bleed was carried out at 14 dpi. Hyperimmune RHDV sera from an additional four rabbits harvested at various time points were available from a previous study (Matthaei et al., 2014). Seven RCV-A1 polyclonal sera were also included (Liu et al., 2012b), as well as an additional seven negative control sera.

For the production of RHDV2 polyclonal sera, rabbits needed to be partially vaccinated prior to the challenge, as young rabbits are not innately resistant to lethal RHDV2 infection (Neimanis et al., 2018a). Three five-week old rabbits and two adult rabbits of approximately three years of age were injected with the experimental vaccine preparation as described above. One adult animal was challenged 10 days after vaccination (dpv) the remaining four rabbits in this group were challenged six dpv. Monitoring and blood sampling of these animals was carried out as for the RHDVa infected rabbits (Table 1).

### RHDV2 specific monoclonal antibody

A mouse monoclonal antibody (Mab) conjugated to horse radish peroxidase (4H12-HRP, lot 150416) and raised against a European strain of RHDV2 was purchased from the Instituto Zooprofilatttico Sperimentale (IZS, Brescia, Italy).

### ELISA

The RHDV IgM, IgA and IgG isotype ELISA (Liu et al., 2012b), the RHDV competition ELISA (cELISA) (Cooke, 2002) and the RCV-A1 blocking ELISA (bELISA) (Liu et al., 2012a) were performed as described previously.

The RHDV2 cELISA workflow and volumes used were similar to the RHDV cELISA, with modifications. Plates were coated overnight at 4°C with a polyclonal RHDV2 rabbit serum (A#Jul, 17 dpi) at a 1:2000 dilution in carbonate buffer (pH 9.6). The plates were washed with PBS –Tween (PBS-T) prior to the addition of serial dilutions of rabbit sera starting at 1:40 dilution followed immediately with the addition of RHDV2 crude antigen at a 1:20 dilution and incubated at 37°C for 1 hour. The plate was washed and the MAb 4H12-HRP at a 1:300 dilution (0.8 µg/ml) was added and incubated at 37°C for 1 hr. The plate was washed and OPD substrate was added and incubated for 5 minutes before being stopped by the addition of 1M sulphuric acid. The plate was read at 492nM. Optimal concentrations of antigen and MAb were determined by checkerboard titrations using rabbit sera known to be positive and negative to RHDV2. Concentrations were selected that resulted in an optical density (OD) of between 1 and 1.2 for a panel of 15 negative reference sera as well as the highest signal to noise ratio when compared to the positive control serum. A serum was scored positive when the OD was reduced to 75% or less compared to the OD of the negative reference serum at the same dilution.

The IgM and IgA ELISAs were adapted to RHDV2 to assess if the sensitivity of detection of these immunoglobulins can be increased. The general set up of the assays was similar to that reported for RHDV IgM and IgA isotype ELISAs (Liu et al., 2012b), except for substituting RHDV virus like particles with crude RHDV2 virus antigen at a 1:20 dilution, and using MAb 4H12-HRP as the detecting antibody at a 1:300 dilution (0.8 µg/ml).

### RHDV2 cELISA analysis of field samples collected before and after RHDV2 arrival

A long term field site located in the central tablelands of New South Wales (Oakey Creek, 33°24’40’’S, 149°22’1’’E) was visited quarterly to collect serum samples from between ten and 25 healthy shot rabbits (average n=19.3), between January 2012 and January 2018. Rabbit shooting and sample collection was carried out as described previously (Cox et al., 2017), rabbit age was estimated based on dry eye lens weight as previously reported (Augusteyn, 2007). Since January 2012, samples collected for this site had been analysed using a suite of ELISAs originally developed for RHDV (Cooke, 2002).

## Results

### Serum production

None of the five week old rabbits infected with RHDVa-Aus or RHDVa-K5 developed severe clinical signs or succumbed to RHD. A short fever episode of ≥40°C was observed in all four kittens infected with RHDVa-Aus and three out of four kittens infected with RHDVa-K5 between 2 and 4 dpi. These fever episodes were not associated with any changes in behavior and all animals continued to gain weight throughout the experiment.

In the group of rabbits used to produce RHDV2 polyclonal serum, no adverse reaction to the experimental vaccine was observed in any of the animals at the injection sites. No behavioral abnormalities or elevated temperatures were recorded in any of the rabbits in this group following vaccination and subsequent challenge with RHDV2.

### Cross-reactivity of reference sera in RHDV isotype ELISAs

Isotype ELISAs, in particular IgM and IgA are important tools to infer disease lagovirus dynamics in rabbit populations, with IgM indicative of a recent outbreak and a boost in IgA indicating re-exposure of previously infected individuals. High levels of cross-reactivity in the RHDV IgM, IgA and IgG isotype ELISAs were observed in sera raised against RHDVa-Aus and RHDVa-K5 compared to the RHDV-Czech polyclonal sera (Table 1). Sera raised against RHDV2 also showed varying levels of cross-reactivity in this assay, but with different patterns. Cross-reactivity was highest in the IgG ELISA, with detectable titres in all animals at 8 dpi, and in one case day 6 post vaccination and prior to RHDV2 challenge. IgM antibodies were detected in all RHDV2 polyclonal reference sera although titres were lower compared to those of the RHDVa-K5, RHDV-Aus and RHDV-Czech reference sera collected at similar time points post infection. IgA antibodies are indicative of active replication of RHDV following a natural infection and are not seen in rabbits treated with inactivated vaccines (Lavazza and Capucci, 2016) and accordingly, no IgA responses were observed in this group following vaccination. IgA cross-reactivity in the RHDV2 polyclonal sera was present but with overall lower titres and with a later onset compared to rabbits infected with RHDV and RHDVa strains, indicating an active but possibly attenuated infection with RHDV2 in these previously vaccinated animals (Table 1). Overall, the IgM and IgA isotype ELISAs originally developed for RHDV appear suitable to detect disease activity of all circulating virulent strains in wild rabbit populations, although the sensitivity of detection may be slightly reduced.

### Increased sensitivity and specificity in the RHDV2 adapted IgA and IgM ELISAs

When the isotype ELISAs for IgA and IgM were adapted to RHDV2 specific reagents, the titres of the RHDV2 reference sera were at least four fold greater when compared to the respective original RHDV isotype assays, and where sufficient amounts of serum were available for testing, both IgM and IgA antibodies were detected earlier (Table 1). RHDV-Czech, RHDVa-K5, and RHDVa-Aus reference sera also cross-reacted in the RHDV2 IgM and IgA ELISA, but had lower titres compared to the RHDV2 reference sera (Table 1). Due to the high level of cross reactivity of RHDV2 polyclonal sera in the in the IgG assay, the IgG isotype ELISA was not adapted.

### Cross-reactivity of reference sera in specific competition or blocking ELISAs for RCV-A1, RHDV and RHDV2

The RHDV2 cELISA developed here from Australian and European reagents proved to be highly sensitive and specific. Low levels of reactivity were detected as early as 6 dpv with titres increasing until the end of the trial (Table 1). Sera raised against RHDV-Czech, RHDVa-Aus and RHDVa-K5 only showed low levels of cross-reactivity in this assay, only in one case exceeding titres of 1:40 (1:160, K#315, 22 dpi, Table 1). None of the seven archival RCV-A1 control sera reacted in the RHDV2 cELISA.

In contrast, the RHDV cELISA showed low to moderate levels of cross-reactivity with RHDV2 polyclonal sera, and very high levels of cross-reactivity with the sera raised against the two RHDVa strains (Table 1). Only one of the RHDV-Czech reference sera tested here showed a very low level of cross-reactivity with the RCV-A1 bELISA (1:10, K#177, 14 dpi, Table 1). Only sera from the terminal bleeds were tested in the RHDV cELISA and RCV-A1 bELISA as there was not sufficient serum left from the previous sampling points.

### The ratio of the two specific cELISAs for RHDV and RHDV2 can be used to infer the presence of RHDV2 specific antibodies

We explored if the ratios between the RHDV and RHDV2 cELISAs could be used to infer the presence of RHDV2 antibodies in the populations, similar to an approach used previously to discriminate between RHDV2 and European Brown Hare Syndrome Virus (EBHSV) antibodies in wild hare populations in Europe (Velarde et al., 2017). While there was some level of cross-reactivity between the respective cELISAs in the reference sera produced for this study, the titres were always higher for the respective specific viral antigen. To investigate this further, negative results were set to a titre of 1:5 for the purpose of forming ratios between reciprocal titres, and only samples were included in the analysis that had returned a positive reaction in either of the two assays, as described previously (Velarde et al., 2017). Due to the high likelihood of cross reactivity at very low dilutions, only test results > 1:40 were considered for this analysis. The RHDV2 cELISA/RHDV cELISA titre ratios were >1 in all reference sera raised against RHDV2, with ratios ranging from 1 to 100. In all reference sera raised against RHDV or RHDVa the ratios were <1, ranging from 0.008 to 0.125 for sera raised against RHDVa-K5 and RHDVa-Aus, and from 0.0002 to 0.0156 for sera raised against RHDV-Czech.

We then applied this method to retrospectively test archival serum samples collected at the Oakey Creek long term study site where these were available. Here, approximately 20 rabbits had been sampled quarterly since January 2012 and analysed with the serological assays originally developed for RHDV (Cooke et al., 2000). RHDV IgM and IgA assays were only carried out at a 1:40 dilution to detect presence or absence of recent disease activity but were not titrated. These samples were re-tested in the RHDV2 cELISA described here to determine the time of RHDV2 arrival at this site. Initially we analysed a very early sample of n=20 rabbits from autumn 2012 as a negative baseline, as it was considered very unlikely that RHDV2 would have been present in Australia over three years prior to its detection. We then continued the analysis starting with the most recent samples, working backwards until four consecutive samples showed no serological evidence of RHDV2. Unexpectedly, serological analysis revealed clear evidence for presence of RHDV2 antibodies in this population with several samples resulting in an RHDV2cELISA/RHDVcELISA ratio of >1 as early as January 2015 (Figure 1). This indicates that RHDV2 must have arrived at this site sometime between the sampling periods October 2014 and January 2015, which pre-dates the first case report of RHDV2 in a deceased rabbit by at least six months (Hall et al., 2015). While fluctuating, overall seroprevalence to RHDV2 increased in later sampling periods coinciding with an overall decrease of animals classified as positive to RHDV.

**Figure 1:**
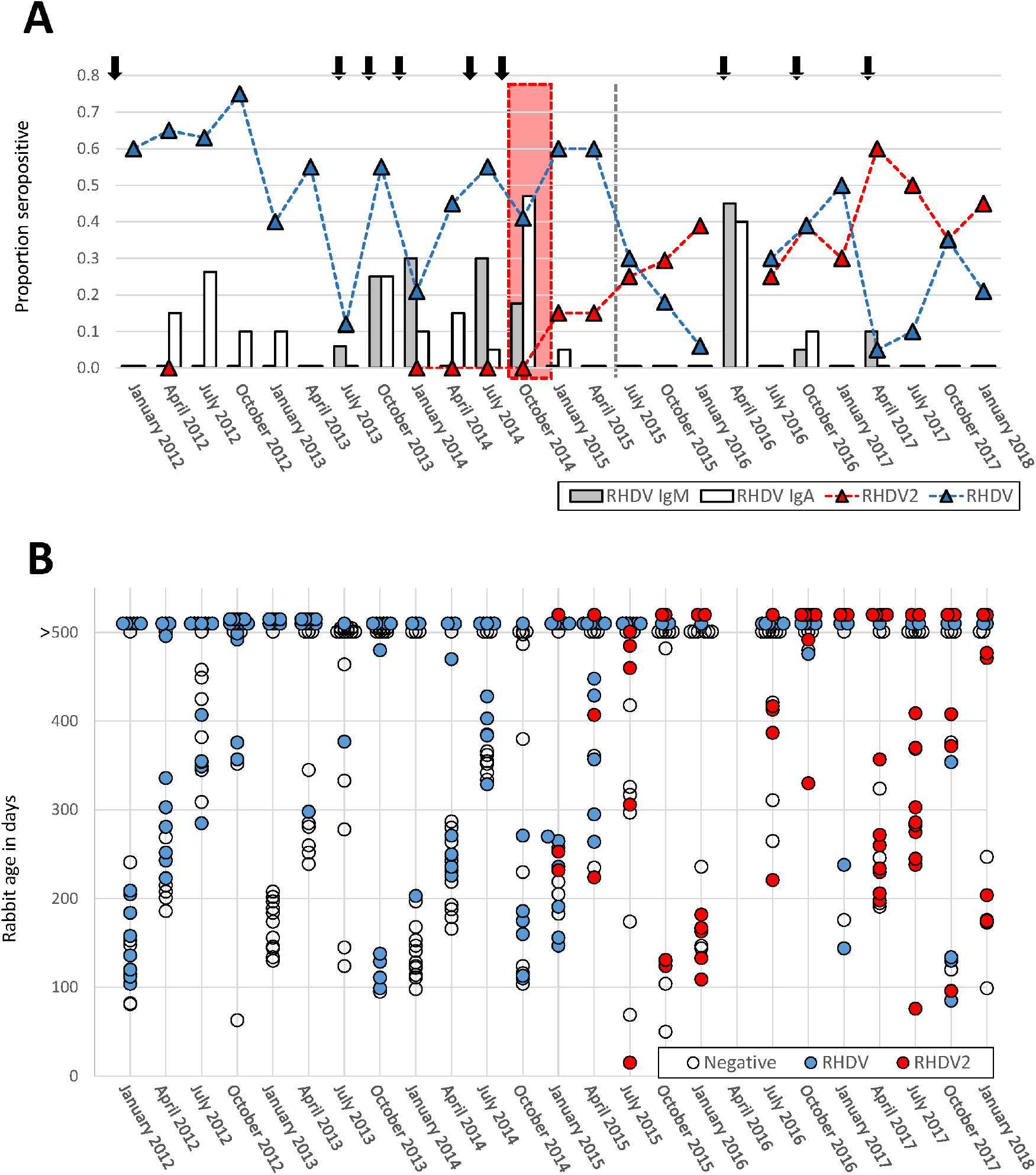
Calicivirus disease activity and proposed arrival of RHDV2 inferred from serological data at a long term monitoring site in NSW (Oakey Creek). A. Sera from between 10 and 25 (avg 19.3) healthy shot rabbits were analysed for RHDV IgM and IgA antibodies four times per year. Presence of IgM antibodies and/or an increase in IgA prevalence is used to infer recent virus activity (Cooke et al., 2000). Rabbits were scored positive to RHDV2 when the ratio of the RHDV2-cELISA/RHDV-cELISA titres was >1 and positive to RHDV when the ratio was <1. Prior to January 2015, RHDV prevalence was inferred as the proportion of rabbits positive to the RHDVcELISA. Black arrows indicate inferred virus outbreaks or prolonged periods of virus activity. The vertical dashed grey line indicates the time RHDV2 was first reported in Australia (Hall et al., 2015). The vertical red box indicates the approximate proposed arrival period of RHDV2 at this site. B. Individual rabbits classified as positive to RHDV or RHDV2 according to their age. Age in days was determined using the dry eye lens weight according to previously described methods (Augusteyn, 2007).

Some level of cross reactivity was observed in the RHDV2 cELISA prior to RHDV2 arrival. Four out of 96 samples analysed that were collected prior to January 2015 showed low level cross reactivity in the RHDV2 cELISA with titres between 1:40 and 1:80. However in all these cases the individual rabbits had higher titres in the RHDV cELISA resulting in a ratio of <1 in all cases.

IgM antibodies indicative of recent lagovirus outbreaks were frequently detected in this population prior to the arrival of RHDV2, namely in winter and spring 2012, summer 2014, and winter and spring 2014. No evidence for recent virus activity was detected in the following five sampling periods, but then again in autumn and spring 2016, and again in autumn 2017 (Figure 1). In order to ascertain which virus was likely causing these outbreaks, we investigated the antibody profiles in the recent cohorts of rabbits (Figure 2). Although some rabbit breeding in this part of Australia is possible year round, the main breeding events occur in winter/spring. Accordingly, members of each new cohort of rabbits starts to appear in the summer shot sample, and subsequent sampling periods throughout the year reflect the increase in age of this cohort (Figure 2). As expected, all animals scored as positive to RHDV with a cELISA ratio <1 prior to January 2015. After the arrival of RHDV2 at this site, in start to appear in the January 2015 sample. However, evidence for RHDV positive animals in less than 12 months of age was found in January 2017 and October 2017. These animals had not yet been born at the time of RHDV2 arrival, indicating that both RHDV and RHDV2 must have been involved in the three periods of virus activity recorded since RHDV2 arrival at this site. The RHDV cELISA titres in these young rabbits were high, ranging from 1:80 to 1:2560 (1:840 avg), indicating a strong immune response from an active RHDV infection rather than the presence of residual maternal antibodies.

## Discussion

Specific serology of different lagoviruses can be challenging, as high levels of cross-reactivity are often observed even between antigenically very different members of this genus (Capucci et al., 1996b, Liu et al., 2012b, Nagesha et al., 2000). For the more specific detection of different viruses, less sensitive and more specific cELISAs or bELISAs have been described (Capucci et al., 1991, Collins et al., 1995, Liu et al., 2012a), however, isotype ELISAs detecting IgG antibodies are so cross-reactive that they have historically been used to infer the presence of previously unknown and often antigenically quite different lagoviruses (Capucci et al., 1996b, Cooke et al., 2000, Nagesha et al., 2000, Robinson et al., 2002).

These difficulties notwithstanding, there is a need for improved serological tools to discriminate between antibodies to the different lagoviruses circulating in Australia, to better understand their role in naturally occurring disease dynamics as well as biological control operations of wild rabbits. For over two decades only RHDV and RCV-A1 were known to be present in Australia, and their disease dynamics have been studied extensively, utilising specific cELISAs or bELISAs for RHDV and RCV-A1 as well as isotype ELISAs for both viruses (Cooke et al., 2000, Liu et al., 2012a, Liu et al., 2012b). The recent arrival of two different strains of RHDVa (RHDVa-Aus and RHDVa-K5) as well as RHDV2 has highlighted the need to update and extend the panel of serological tools available on this continent to enable better studies of the epidemiology and interactions of these viruses.

Disease dynamics of RHDV can be inferred by interpreting the reaction profiles in different isotype and cELISAs (Cooke, 2002). In this approach, IgM is interpreted as an indicator of a recent outbreak and first time exposure of an individual to RHDV, a boost in IgA titres as a measure for re-exposure to RHDV, and the highly sensitive but cross-reactive IgG is used to infer the presence of maternal antibodies in very young rabbits. In our study, the reference sera raised against RHDV-Czech, RHDVa-Aus, RHDVa-K5 and RHDV2 all reacted to varying degrees in these assays, indicating that they should be suitable to detect broad patterns of disease activity caused by any of the virulent lagovirus strains in wild rabbit populations in Australia, however they do not allow for the discrimination between the various strains.

Adaptation of the IgM and IgA ELISA to RHDV2 by using RHDV2-specific antigen and Mab increased the sensitivity of these assays for RHDV2 IgM and IgA detection substantially. While the use of the original RHDV IgA and IgM assays is sufficient to infer broader disease activity patterns in wild rabbit populations for all lagoviruses including RHDV2 (Figure 1), switching to these more sensitive assays for future large scale field epidemiology studies should be considered if RHDV2 remains the dominant strain in the Australian landscape (Mahar et al., 2018a).

The RHDV2 cELISA described here proved to be highly specific for the detection of RHDV2 antibodies, with low levels of cross-reactivity to the experimentally produced RHDV and RHDVa reference sera as well as the sera from wild rabbits collected before the arrival of RHDV2 in Australia. In contrast, the existing RHDV cELISA (Capucci et al., 1991) showed low to medium levels of cross-reactivity to RHDV2 and high levels of cross-reactivity to both RHDVa-Aus and RHDVa-K5. However, in the RHDV and RHDV2 cELISAs, titres were higher in the sera raised against the respective strain, such that the ratios of the reciprocal titres of the two assays can be used to infer which strain the rabbit was most likely exposed to. It needs to be noted that this method classifies rabbits into positive to either RHDV or RHDV2, and therefore does not allow for the detection of mixed infections and the true prevalence to both RHDV and RHDV2 is likely underestimated. However the method does allow to detect presence of RHDV2 and should allow to discern broader trends within rabbit populations.

When applied to a sample collection from a long term field monitoring site, this method showed clear evidence for RHDV2 activity in this population at least six months prior to the first RHDV2 case reported in Australia (Hall et al., 2015), and confirms inferences made from phylogenetic analyses of viral sequences suggesting that RHDV2 had circulated in Australia several months prior to its first detection in May 2015 (Mahar et al., 2018a).

The inclusion of age data into the analysis allowed the confirmation of RHDV activity in this population after July 2016. In addition, including age data into the analysis may provide a more accurate estimate of disease dynamics than analysing overall seroprevalence data alone. A large proportion of rabbits present in every shot sample were >500 days old, and may therefore confound the analysis of recent virus activity. In addition, if some level of immunological cross protection exists between RHDV and RHDV2 (Calvete et al., 2018), the removal of susceptible rabbits by an RHDV2 outbreak from the population could result in an apparent increase of RHDV seroprevalence, due to the resulting increased proportion of surviving old RHDV seropositive animals in the sample.

## Conclusions

The analysis of the reference sera as well as field sera collected pre- and post-RHDV2 arrival in Australia indicates that existing RHDV IgA and IgM isotype ELISAs are suitable to infer disease dynamics of all virulent RHDV strains currently circulating in Australia. The ratio between the RHDV2 and RHDV cELISA titres allows for the detection of the presence of RHDV2 specific antibodies with high levels of certainty, even in the presence of RHDV or RHDVa antibodies. In combination, the methods described here should allow for retrospective analysis of archival field sera to study the spread and impact of RHDV2 on Australian rabbit populations as well as allow for a more accurate estimate of the time of RHDV2 arrival at various site in Australia. Including rabbit age data, where available, will improve these analyses. However, due to the high levels of cross-reactivity of RHDV2 reference sera in the RHDV cELISA, detecting RHDV or RHDVa antibodies in samples collected post RHDV2 arrival is challenging. In particular, inferring mixed infections of RHDV/RHDVa/RHDV2 or confirming the absence of RHDV antibodies in RHDV2 positive populations is difficult to discern. Similarly, distinguishing between antibodies to RHDV and RHDVa strains is not feasible with the serological tools currently available. Despite these remaining challenges in the differential serodiagnostics of Australian lagoviruses, the additional assays described here represent an important addition to the tool kit that will benefit ongoing continent-wide lagovirus epidemiology studies.

## Acknowledgements

We thank Peter Kerr and Robyn Hall for helpful comments on the manuscript.

## References

Augusteyn, R. C., 2007: On the relationship between rabbit age and lens dry weight: improved determination of the age of rabbits in the wild. Mol Vis, 13, 2030–2034.

Barcena, J., B. Guerra, I. Angulo, J. Gonzalez, F. Valcarcel, C. P. Mata, J. R. Caston, E. Blanco and A. Alejo, 2015: Comparative analysis of rabbit hemorrhagic disease virus (RHDV) and new RHDV2 virus antigenicity, using specific virus-like particles. Veterinary research, 46, 106.

Calvete, C., M. Mendoza, A. Alcaraz, M. P. Sarto, M. P. Jimenez-de-Baguess, A. J. Calvo, F. Monroy and J. H. Calvo, 2018: Rabbit haemorrhagic disease: Cross-protection and comparative pathogenicity of GI.2/RHDV2/b and GI.1b/RHDV lagoviruses in a challenge trial. Vet Microbiol, 219, 87–95.

Camarda, A., N. Pugliese, P. Cavadini, E. Circella, L. Capucci, A. Caroli, M. Legretto, E. Mallia and A. Lavazza, 2014: Detection of the new emerging rabbit haemorrhagic disease type 2 virus (RHDV2) in Sicily from rabbit (Oryctolagus cuniculus) and Italian hare (Lepus corsicanus). Research in Veterinary Science, 97, 642–645.

Capucci, L., D. Chasey, A. Lavazza and D. Westcott, 1996a: Preliminary characterization of a non-haemagglutinating strain of rabbit haemorrhagic disease virus from the United Kingdom. Zentralbl Veterinarmed B, 43, 245–250.

Capucci, L., F. Fallacara, S. Grazioli, A. Lavazza, M. L. Pacciarini and E. Brocchi, 1998: A further step in the evolution of rabbit hemorrhagic disease virus: the appearance of the first consistent antigenic variant. Virus Res, 58, 115–126.

Capucci, L., G. Frigoli, L. Ronshold, A. Lavazza, E. Brocchi and C. Rossi, 1995: Antigenicity of the rabbit hemorrhagic disease virus studied by its reactivity with monoclonal antibodies. Virus Res, 37, 221–238.

Capucci, L., P. Fusi, A. Lavazza, M. L. Pacciarini and C. Rossi, 1996b: Detection and preliminary characterization of a new rabbit calicivirus related to rabbit hemorrhagic disease virus but nonpathogenic. J Virol, 70, 8614–8623.

Capucci, L., M. T. Scicluna and A. Lavazza, 1991: Diagnosis of viral haemorrhagic disease of rabbits and the European brown hare syndrome. Rev Sci Tech, 10, 347–370.

Carvalho, C. L., E. L. Duarte, M. Monteiro, A. Botelho, T. Albuquerque, M. Fevereiro, A. M. Henriques, S. S. Barros and M. D. Duarte, 2017: Challenges in the rabbit haemorrhagic disease 2 (RHDV2) molecular diagnosis of vaccinated rabbits. Vet Microbiol, 198, 43–50.

Collins, B. J., J. R. White, C. Lenghaus, V. Boyd and H. A. Westbury, 1995: A Competition Elisa For The Detection Of Antibodies To Rabbit Hemorrhagic-Disease Virus. Vet Microbiol, 43, 85–96.

Cooke, B., Chudleigh, P., Simpson S., Saunders, G., 2013: The economic Benefits of the biological control of rabbits in Australia, 150 - 2011. Australian Economic History Review, 53, 1–17.

Cooke, B. D., 2002: Rabbit haemorrhagic disease: field epidemiology and the management of wild rabbit populations. Rev Sci Tech, 21, 347–358.

Cooke, B. D., R. P. Duncan, I. McDonald, J. Liu, L. Capucci, G. J. Mutze and T. Strive, 2018: Prior exposure to non-pathogenic calicivirus RCV-A1 reduces both infection rate and mortality from rabbit haemorrhagic disease in a population of wild rabbits in Australia. Transboundary and emerging diseases, 65, e470–e477.

Cooke, B. D. and F. Fenner, 2002: Rabbit haemorrhagic disease and the biological control of wild rabbits, Oryctolagus cuniculus, in Australia and New Zealand. Wildlife Research, 29, 689–706.

Cooke, B. D., A. J. Robinson, J. C. Merchant, A. Nardin and L. Capucci, 2000: Use of ELISAs in field studies of rabbit haemorrhagic disease (RHD) in Australia. Epidemiology and infection, 124, 563–576.

Cox, T., T. Strive, G. Mutze, P. West and G. Saunders, 2013: Benefits of Rabbit Control in Australia: Report. Invasive Animals Cooperative research Centre.

Cox, T. E., J. Liu, R. van de Ven and T. Strive, 2017: Different Serological Profiles to Co-Occurring Pathogenic and Nonpathogenic Caliciviruses in Wild European Rabbits (Oryctolagus Cuniculus) across Australia. J Wildlife Dis, 53, 472–481.

Dalton, K. P., I. Nicieza, J. Abrantes, P. J. Esteves and F. Parra, 2014: Spread of new variant RHDV in domestic rabbits on the Iberian Peninsula. Vet Microbiol, 169, 67–73.

Dalton, K. P., I. Nicieza, A. Balseiro, M. A. Muguerza, J. M. Rosell, R. Casais, A. L. Alvarez and F. Parra, 2012: Variant rabbit hemorrhagic disease virus in young rabbits, Spain. Emerg Infect Dis, 18, 2009–2012.

Dalton, K. P., A. Podadera, V. Granda, I. Nicieza, D. del Llano, R. Gonzalez, J. R. de Los Toyos, M. G. Ocana, F. Vazquez, J. M. M. Alonso, J. M. Prieto, F. Parra and R. Casais, 2018: ELISA for detection of variant rabbit haemorrhagic disease virus RHDV2 antigen in liver extracts. J Virol Methods, 251, 38–42.

Duarte, M. D., C. L. Carvalho, S. C. Barros, A. M. Henriques, F. Ramos, T. Fagulha, T. Luis, E. L. Duarte and M. Fevereiro, 2015: A real time Taqman RT-PCR for the detection of rabbit hemorrhagic disease virus 2 (RHDV2). J Virol Methods, 219, 90–95.

Elsworth, P. G., J. Kovaliski and B. D. Cooke, 2012: Rabbit haemorrhagic disease: are Australian rabbits (Oryctolagus cuniculus) evolving resistance to infection with Czech CAPM 351 RHDV? Epidemiology and infection, 140, 1972–1981.

Hall, R. N., J. E. Mahar, S. Haboury, V. Stevens, E. C. Holmes and T. Strive, 2015: Emerging Rabbit Hemorrhagic Disease Virus 2 (RHDVb), Australia. Emerg Infect Dis, 21, 2276–2278.

Hall, R. N., J. E. Mahar, A. J. Read, R. Mourant, M. Piper, N. Huang and T. Strive, 2018: A strain-specific multiplex RT-PCR for Australian rabbit haemorrhagic disease viruses uncovers a new recombinant virus variant in rabbits and hares. Transboundary and emerging diseases, 65, e444–e456.

Hall, R. N., D. E. Peacock, J. Kovaliski, J. E. Mahar, R. Mourant, M. Piper and T. Strive, 2017: Detection of RHDV2 in European brown hares (Lepus europaeus) in Australia. Vet Rec, 180.

Kong, D. S., J. S. Liu, Q. Jiang, Z. Yu, X. L. Hu, D. C. Guo, Q. Q. Huang, M. H. Jiao and L. D. Qu, 2016: Production, Characterization, and Epitope Mapping of Monoclonal Antibodies Against Different Subtypes of Rabbit Hemorrhagic Disease Virus (RHDV). Sci Rep-Uk, 6.

Lavazza, A. and L. Capucci, 2016: Rabbit haemorrhagic disease. In: OIE manual of diagnostic tests and vaccines for terrestrial animals. 2016 [cited 2017 Jan 13]. http://www.oie.int/fileadmin/Home/eng/Health_standards/tahm/2.06.02_RHD.pdf.

Le Gall-Recule, G., E. Lemaitre, S. Bertagnoli, C. Hubert, S. Top, A. Decors, S. Marchandeau and J. S. Guitton, 2017: Large-scale lagovirus disease outbreaks in European brown hares (Lepus europaeus) in France caused by RHDV2 strains spatially shared with rabbits (Oryctolagus cuniculus). Veterinary research, 48.

Le Gall-Recule, G., F. Zwingelstein, S. Boucher, B. Le Normand, G. Plassiart, Y. Portejoie, A. Decors, S. Bertagnoli, J. L. Guerin and S. Marchandeau, 2011: Detection of a new variant of rabbit haemorrhagic disease virus in France. Vet Rec, 168, 137–138.

Le Pendu, J., J. Abrantes, S. Bertagnoli, J. S. Guitton, G. Le Gall-Recule, A. M. Lopes, S. Marchandeau, F. Alda, T. Almeida, A. P. Celio, J. Barcena, G. Burmakina, E. Blanco, C. Calvete, P. Cavadini, B. Cooke, K. Dalton, M. D. Mateos, W. Deptula, J. S. Eden, F. Wang, C. C. Ferreira, P. Ferreira, P. Foronda, D. Goncalves, D. Gavier-Widen, R. Hall, B. Hukowska-Szematowicz, P. Kerr, J. Kovaliski, A. Lavazza, J. Mahar, A. Malogolovkin, R. M. Marques, S. Marques, A. Martin-Alonso, P. Monterroso, S. Moreno, G. Mutze, A. Neimanis, P. Niedzwiedzka-Rystwej, D. Peacock, F. Parra, M. Rocchi, C. Rouco, N. Ruvoen-Clouet, E. Silva, D. Silverio, T. Strive, G. Thompson, B. Tokarz-Deptula and P. Esteves, 2017: Proposal for a unified classification system and nomenclature of lagoviruses. Journal of General Virology, 98, 1658–1666.

Liu, J., D. A. Fordham, B. D. Cooke, T. Cox, G. Mutze and T. Strive, 2014: Distribution and Prevalence of the Australian Non-Pathogenic Rabbit Calicivirus Is Correlated with Rainfall and Temperature. Plos One, 9.

Liu, J., P. J. Kerr and T. Strive, 2012a: A sensitive and specific blocking ELISA for the detection of rabbit calicivirus RCV-A1 antibodies. Virol J, 9, (3 September 2012)-(2013 September 2012).

Liu, J., P. J. Kerr, J. D. Wright and T. Strive, 2012b: Serological assays to discriminate rabbit haemorrhagic disease virus from Australian non-pathogenic rabbit calicivirus. Vet Microbiol, 157, 345–354.

Mahar, J. E., R. N. Hall, D. Peacock, J. Kovaliski, M. Piper, R. Mourant, N. N. Huang, S. Campbell, X. N. Gu, A. Read, N. Urakova, T. Cox, E. C. Holmes and T. Strive, 2018a: Rabbit Hemorrhagic Disease Virus 2 (RHDV2; GI.2) Is Replacing Endemic Strains of RHDV in the Australian Landscape within 18 Months of Its Arrival. J Virol, 92.

Mahar, J. E., A. J. Read, X. N. Gu, N. Urakova, R. Mourant, M. Piper, S. Haboury, E. C. Holmes, T. Strive and R. N. Hall, 2018b: Detection and Circulation of a Novel Rabbit Hemorrhagic Disease Virus in Australia. Emerg Infect Dis, 24, 22–31.

Matthaei, M., P. J. Kerr, A. J. Read, P. Hick, S. Haboury, J. D. Wright and T. Strive, 2014: Comparative quantitative monitoring of rabbit haemorrhagic disease viruses in rabbit kittens. Virol J, 11.

Mutze, G., P. Bird, S. Jennings, D. Peacock, N. de Preu, J. Kovaliski, B. Cooke and L. Capucci, 2014: Recovery of South Australian rabbit populations from the impact of rabbit haemorrhagic disease. Wildlife Research, 41, 552–559.

Mutze, G., B. Cooke and P. Alexander, 1998: The initial impact of rabbit hemorrhagic disease on European rabbit populations in South Australia. J Wildl Dis, 34, 221–227.

Nagesha, H. S., K. A. McColl, B. J. Collins, C. J. Morrissy, L. F. Wang and H. A. Westbury, 2000: The presence of cross-reactive antibodies to rabbit haemorrhagic disease virus in Australian wild rabbits prior to the escape of virus from quarantine. Arch Virol, 145, 749–757.

Neave, M. J., R. N. Hall, N. Huang, K. A. McColl, P. Kerr, M. Hoehn, J. Taylor and T. Strive, 2018: Robust Innate Immunity of Young Rabbits Mediates Resistance to Rabbit Hemorrhagic Disease Caused by Lagovirus Europaeus GI.1 But Not GI.2. Viruses, 10.

Neimanis, A., U. L. Pettersson, N. Huang, D. Gavier-Widen and T. Strive, 2018a: Elucidation of the pathology and tissue distribution of Lagovirus europaeus GI.2/RHDV2 (rabbit haemorrhagic disease virus 2) in young and adult rabbits (Oryctolagus cuniculus). Veterinary research, 49.

Neimanis, A. S., H. Ahola, U. L. Pettersson, A. M. Lopes, J. Abrantes, S. Zohari, P. J. Esteves and D. Gavier-Widen, 2018b: Overcoming species barriers: an outbreak of Lagovirus europaeus GI.2/RHDV2 in an isolated population of mountain hares (Lepus timidus). Bmc Vet Res, 14.

Nystroem, K., G. Le Gall-Recule, P. Grassi, J. Abrantes, N. Ruvoen-Clouet, B. Le Moullac-Vaidye, A. M. Lopes, P. J. Esteves, T. Strive, S. Marchandeau, A. Dell, S. M. Haslam and J. Le Pendu, 2011: Histo-Blood Group Antigens Act as Attachment Factors of Rabbit Hemorrhagic Disease Virus Infection in a Virus Strain-Dependent Manner. Plos Pathog, 7.

Oem, J. K., K. N. Lee, I. S. Roh, K. K. Lee, S. H. Kim, H. R. Kim, C. K. Park and Y. S. Joo, 2009: Identification and Characterization of Rabbit Hemorrhagic Disease Virus Genetic Variants Isolated in Korea. Journal of Veterinary Medical Science, 71, 1519–1523.

Pedler, R. D., R. Brandle, J. L. Read, R. Southgate, P. Bird and K. E. Moseby, 2016: Rabbit biocontrol and landscape-scale recovery of threatened desert mammals. Conservation Biology.

Puggioni, G., P. Cavadini, C. Maestrale, R. Scivoli, G. Botti, L. Ciriaco, G. Le Gall-Recule, A. Lavazza and L. Capucci, 2013: The new French 2010 Rabbit Hemorrhagic Disease Virus causes an RHD-like disease in the Sardinian Cape hare (Lepus capensis mediterraneus). Veterinary research, 44, 1–14.

Robinson, A. J., P. D. Kirkland, R. I. Forrester, L. Capucci, B. D. Cooke and A. W. Philbey, 2002: Serological evidence for the presence of a calicivirus in Australian wild rabbits, Oryctolagus cuniculus, before the introduction of rabbit haemorrhagic disease virus (RHDV): its potential influence on the specificity of a competitive ELISA for RHDV. Wildlife Research, 29, 655–662.

Saunders, G., B. Kay, G. Mutze and D. Choquenot, 2002: Observations on the impacts of rabbit haemorrhagic disease on agricultural production values in Australia. Wildlife Research, 29, 605–613.

Strive, T. and T. E. Cox, 2019: Lethal biological control of rabbits–the most powerful tools for landscape-scale mitigation of rabbit impacts in Australia. Australian Zoologist.

Strive, T., P. Elsworth, J. Liu, J. D. Wright, J. Kovaliski and L. Capucci, 2013: The non-pathogenic Australian rabbit calicivirus RCV-A1 provides temporal and partial cross protection to lethal Rabbit Haemorrhagic Disease Virus infection which is not dependent on antibody titres. Veterinary research, 44, (8 July 2013)-(2018 July 2013).

Velarde, R., P. Cavadini, A. Neimanis, O. Cabezon, M. Chiari, A. Gaffuri, S. Lavin, G. Grilli, D. Gavier-Widen, A. Lavazza and L. Capucci, 2016: Spillover Events of Infection of Brown Hares (Lepus europaeus) with Rabbit Haemorrhagic Disease Type 2 Virus (RHDV2) Caused Sporadic Cases of an European Brown Hare Syndrome-Like Disease in Italy and Spain. Transboundary and emerging diseases.

Velarde, R., P. Cavadini, A. Neimanis, O. Cabezon, M. Chiari, A. Gaffuri, S. Lavin, G. Grilli, D. Gavier-Widen, A. Lavazza and L. Capucci, 2017: Spillover Events of Infection of Brown Hares (Lepus europaeus) with Rabbit Haemorrhagic Disease Type 2 Virus (RHDV2) Caused Sporadic Cases of an European Brown Hare Syndrome-Like Disease in Italy and Spain. Transboundary and emerging diseases, 64, 1750–1761.

Wang, X., H. Hao, L. Qiu, R. Dang, E. Du, S. Zhang and Z. Yang, 2012: Phylogenetic analysis of rabbit hemorrhagic disease virus in China and the antigenic variation of new strains. Arch Virol, 157, 1523–1530.

